# Repression of YdaS Toxin is Mediated by Transcriptional Repressor RacR in the cryptic rac prophage of Escherichia coli-K12

**DOI:** 10.1101/100248

**Authors:** Revathy Krishnamurthi, Swagatha Ghosh, Supriya Khedkar, Aswin Sai Narain Seshasayee

## Abstract

Horizontal gene transfer is a major driving force behind the genomic diversity seen in prokaryotes. The *rac* prophage in *E.coli* K12 encodes a putative transcription factor RacR, whose deletion is lethal. We have shown that the essentiality of *racR* in *E.coli* K12 is attributed to its role in transcriptionally repressing a toxin gene called *ydaS*, which is coded adjacent and divergently to *racR*.

## Introduction

Horizontal Gene Transfer (HGT) contributes to the vast genome diversity seen in prokaryotes. The size of the genomes of *E.coli* varies from 3.97 Mb to 5.85 Mb. The core genome constitutes only ~10% of the gene families represented across these *E.coli* genomes. The rest of the genetic content is variable across strains and often found in genomic islands [^1^]. Many virulence factors and determinants of antibiotic resistance are known to be horizontally acquired, and encoded for example in autonomously replicating plasmids and chromosomally replicating prophages [^2^] [^3^].

The genome of the laboratory strain *E.coli*-K12 comprises nine cryptic prophages which constitute 3.6% of its total genome. The successful maintenance of any horizontally acquired element depends on the conventional selection advantage that it provides to the host and as well as addiction imposed on the host by selfish genetic modules.

Conventional selection is defined by the benefit it provides the host under the given condition. A horizontally acquired gene may integrate into the rest of the cellular network by affecting the function of genes belonging to the core genome, or that of unrelated horizontally acquired elements[^4^][^5^]. For example, a phenotypic microarray study has shown that the nine cryptic prophages in *E.coli*-K12 help the bacterium survive under various stresses [^6^].

Several horizontally acquired elements also carry addiction molecules, including Restriction Modification (R-M) and Toxin Antitoxin (T-A) systems [^7^][^8^]. Loss of such modules can result in post-segregational killing, which encourages the maintenance of such DNA [^9^]. Some T-A systems and R-M systems may have eventually evolved functions which provide benefit to the host, including roles in programmed cell death [^10^][^11^].

During an effort towards addressing the regulatory roles of horizontally acquired transcriptional regulators, we learnt that the poorly characterized gene *racR* which is a putative repressor of the *rac* prophage is an essential gene in *E.coli* K12. Using a combination of genetics, biochemistry and bioinformatics, we present evidence that RacR is indeed a transcriptional repressor. We have shown that RacR binds to its own regulatory region. The adjacent and divergently coded *ydaS* and *ydaT* together encode a toxin whose expression is repressed by the function of RacR. Thus *ydaST* – *racR* module forms a toxin-repressor combination, which makes this RacR regulator essential to the cell.

## Results

### RacR is an essential transcriptional regulator

The *rac* prophage is a cryptic prophage found in *E.coli*. It is a mosaic prophage. It is 23 kb long, and encodes 29 genes in *E.coli* K12. However, its size and gene content vary across *E.coli* strains, with only a few highly conserved genes, which include *recE* – involved in alternative homologous recombination pathway, and *trkG*, a potassium ion permease [Figure 1A].

**Figure1:**
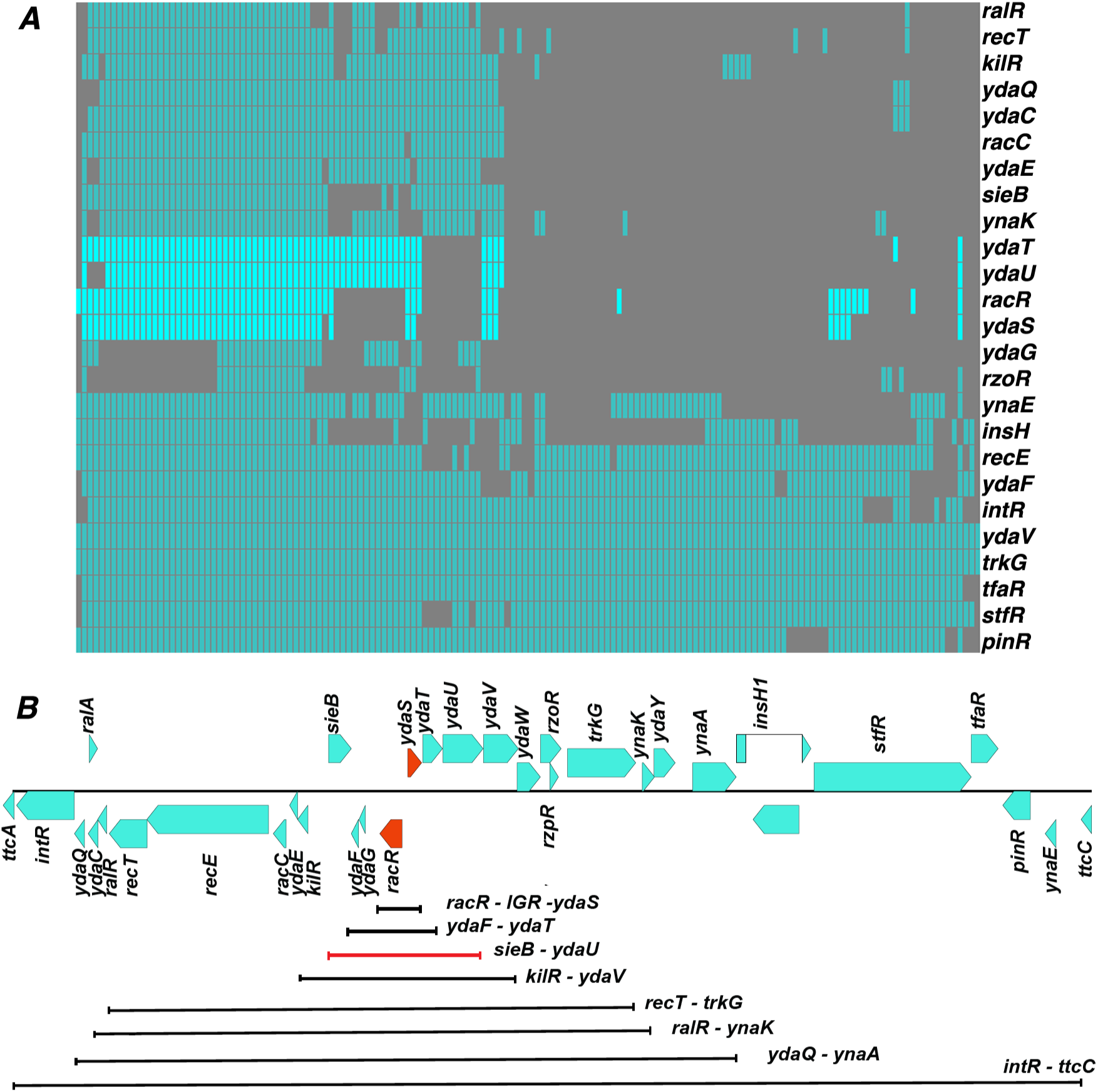
*A) Matrix showing the conservation of rac prophage genes across 154 E.coli genomes. Absence of any gene is indicated by grey and the presence by cyan. The genes racR, ydaS, ydaT and ydaU are shown in bright cyan. Note that the presence of ydaS is always accompanied by the presence of racR. B)Map of rac prophage showing the putative regulatory genes in orange. The lines indicates the eight racR inclusive regions deleted. The line marked in red shows the region deleted from sieB-ydaU which left the gene kilR in the absence of racR.*

Among the less conserved portion of the *rac* prophage is a predicted transcription factor called RacR. It contains a weak helix-turn-helix motif and at best is very distantly related to the lambda *cl* repressor (15% identity by Needleman-Wunsch global alignment). Its deletion is presumed to be lethal. The Keio collection of *E.coli* single gene deletion mutants does not contain *ΔracR* [^12^], and we were unable to delete *racR* by homologous recombination. Nevertheless, the entire *rac* prophage could be deleted (we refer to this as *Δrac* here), and the prophage excises at high rates in certain genetic backgrounds [^13^]. Hence we hypothesized that RacR could be a repressor of a toxin in the same prophage. Because the *rac* prophage carries a previously reported toxin called KilR - an inhibitor of cell division [^14^] - we initiated our screen for the toxin by attempting to delete *racR* in the *ΔkilR* strain. However, we found that *ΔracR* could not be obtained even in a *ΔkilR* background.

We then deleted successively shorter racR-inclusive segments of the prophage. If a deletion attempt removed *racR* but not the toxin that RacR might repress, we would not recover the mutant. The smallest deletion we obtained by this approach included *racR*, its neighboring, divergent gene *ydaS* and the common intergenic region (henceforth referred to as IGR) between them [Figure 1B]. Thus the absence of *ydaS* and the common IGR between *racR* and *ydaS* is a suppressor of the lethality of *ΔracR*.

Despite several attempts, we were unable to delete *racR* in *ΔydaS* without disturbing the IGR between them. However, we obtained *ΔracR* with its IGR intact in a *ΔydaS-T* background. *ydaT* is encoded in tandem and downstream of *ydaS* and might be part of the same operon.

### Over expression of *ydaS* and *ydaS-T* reduces growth

We tested the toxicity of *ydaS*, *ydaT* and *ydaS-T* by cloning these genes under the *araBAD* promoter in pBAD18. Expression of these cloned genes was induced in both wildtype and *Δrac* with 0.1% L-arabinose. We found that the expression of *ydaS* and *ydaS-T* causes rapid growth inhibition after induction in both wildtype and *Δrac* [Supplementary Figure S1]. We collected samples at 5 hours and 14 hours after induction and spotted these on agar plates. Cells expressing *ydaS* and *ydaS-T* from pBAD18 did not grow on these plates [Figure 2A]. The expression of *ydaT* alone did not have any inhibitory or lethal effect on the wild type or the *Δrac* prophage strain.

**Figure2:**
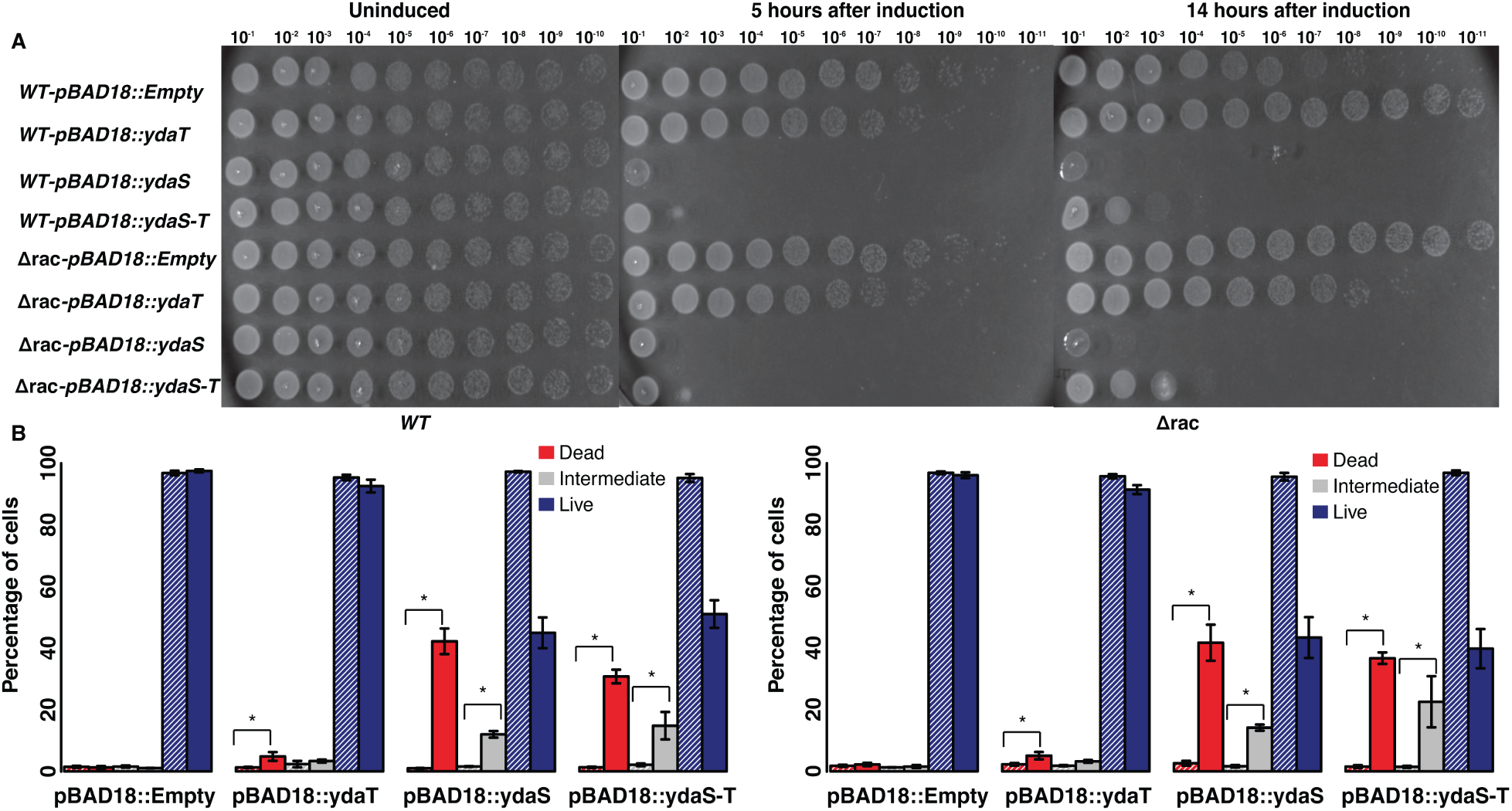
*A) Log and stationary phase cultures of pBAD18-ydaS, pBAD18-ydaT, pBAD18-ydaS-T, and Empty vector in wild type and in Δrac background, grown in the presence or absence of 0.2% L-Arabinose, were spotted on the LB plate without arabinose. B) Live Dead Assay of pBAD18-ydaS, pBAD18-ydaT, pBAD18-ydaS-T and Empty vector in wild type and in Δrac backgrounds. The cells were collected after 5 hours of induction and treated with Propidium Iodide (PI) to mark the dead cells. Bar graph represents the percentage of dead, intermediate and live population of cells in uninduced (striped) and induced (filled)cultures. * corresponds to p-value < 0.01; Wilcoxon rank sum test between induced and uninduced constructs. Error bar here represents the standard error computed from six independent trials (Three biological and two technical replicates).*

Further, we quantified the live and dead cell populations after the induction of *ydaS*, *ydaT* and *ydaS-T* by FACS using Propidium Iodide (PI) as the marker for dead cells. Results from six independent trials show that *ydaS* and *ydaS-T* expression, irrespective of the strain background, leads to loss of cell viability [Figure 2B]. We noticed that *ydaS-T* expressing cells were lengthier than the *ydaS* or *ydaT* expressing cells [Supplementary Figure S2].

Together, these results show that the expression of *ydaS* and *ydaS-T* is lethal, and *ydaS* and *ydaT* do not form a TA pair as predicted earlier [^15^]. YdaS is critical to cell killing, and YdaT may enhance the lethal effect of YdaS while not being toxic on its own.

### Co occurrence of *racR* and *ydaS* implies interaction between them

Functionally related genes tend to be conserved together across genomes [^16^]. We examined the conservation of genes of the *rac* prophage across 154 *E.coli* genomes. Bi-directional best hit search for orthologs confirmed the mosaic nature of the *rac* prophage. In fact, more than 50% of the strains have lost half of the prophage. The genes that are well conserved across the genomes are those, such as *recE* and *trkG*, which have documented functions in the host. Some classical phage genes like *intR*, *pinR*, *stfR*, *tfaR*, *ydaF* and *ydaV* are conserved in more than 85% of the strains analyzed.

We observe that the known toxin genes in the prophage are lost in most of the strains and where present, are always accompanied by its cognate antitoxin genes. RalR-RalA is a known type I T-A system in the same prophage[^17^]. We observe that the RalR toxin is conserved only in 36.3% of the strains we analyzed; the corresponding non-coding antitoxin gene was found in all these strains. KilR, previously reported as a FtsZ inhibitor, was found in 48% of the strains in this analysis; its antitoxin, if any, is unknown.

YdaS is present only in 33.7% of the strains analyzed and we observe that it always co-occurs with RacR [Figure 1A]. A few strains encoded *ydaT* gene in the absence of *racR*; however the IGR was lost in these strains, and certain point mutations were found in the *ydaT* gene. We also observe that YdaT expression on its own, in the absence of YdaS, is not lethal. Thus, genome context analysis suggests a functional interaction between RacR and YdaS(-T).

### Expression of *ydaS* is kept silent under normal physiological conditions

In order to examine the expression of RacR and YdaS in vivo, we tagged these two genes with C-terminal 3X-FLAG (DYKDDDDK). Western blotting using an anti-FLAG antibody showed that RacR was expressed in all the phases of growth. However, YdaS expression could not be detected in our experimental conditions [Fig 3A]. An absolute protein quantification study reported by Li *et.al* 2014., also shows low copy number for YdaS [^18^].

**Figure 3:**
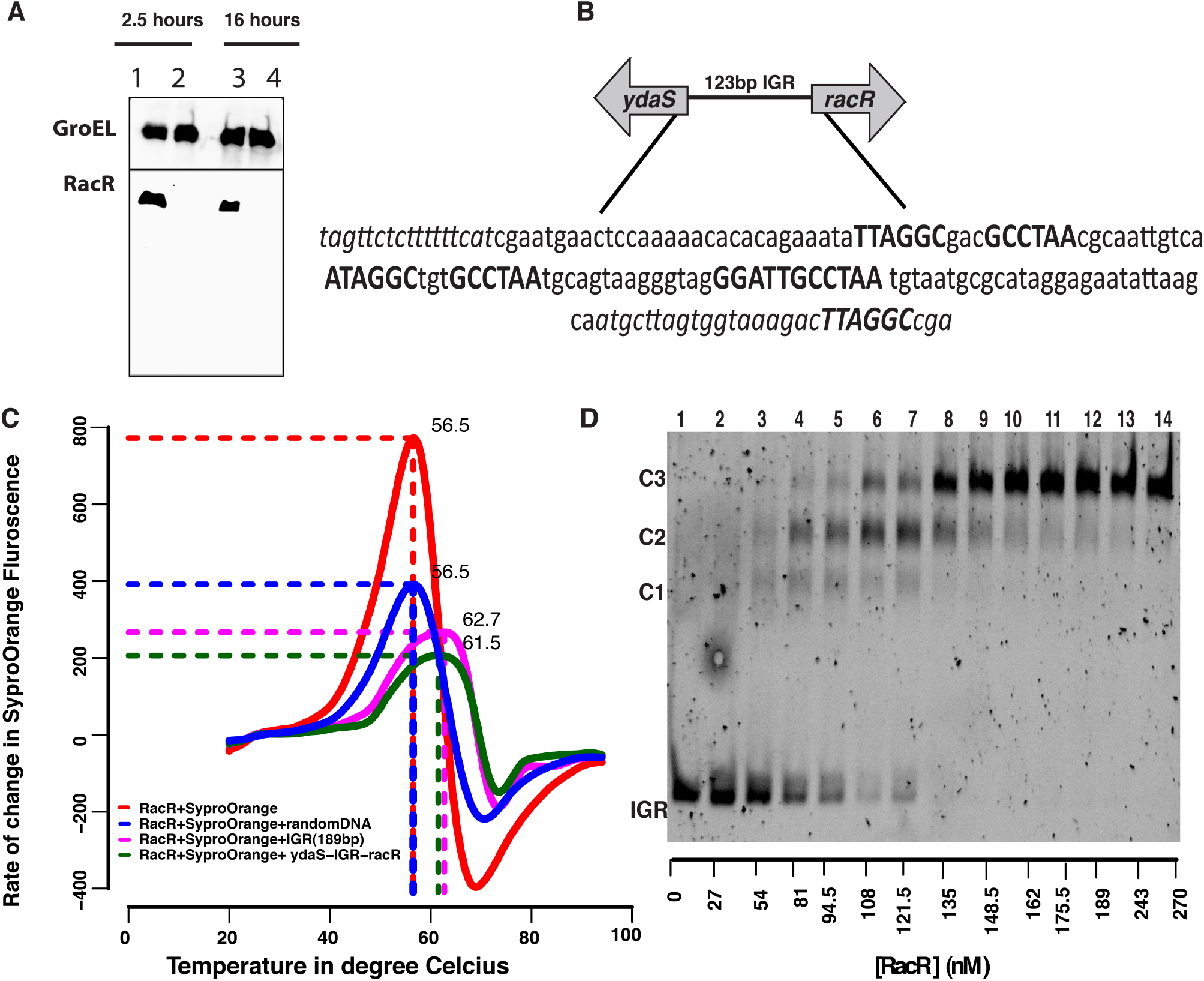
*A)Western blot showing the expression of RacR; YdaS expression could not be detected. The top panel showing GroEL as loading control and the bottom panel showing the RacR expression during log and stationary phase.20μg of total protein was used. B)Intergenic Region (IGR) between ydaS and racR showing the repeat elements with slightly varying sequence in three different regions (bold sequence).Note that the ydaS and racR are coded divergently and in opposite strands. C)Thermal Shift Assay showing ~5°C shift in the TD of RacR in the presence of IGR. D) Non-radioactive Electrophoretic Mobility Shift Assay showing the binding of RacR with IGR with distinct complexes marked as C1,C2 and C3.15 nM of 123bp IGR was titrated against increasing concentration of RacR from 27 nM to 270 nM (Lane 2-14). Lane 1 shows only 15nM of IGR without RacR.*

Analysis of various publicly available and in-house RNA-seq data showed that the expression of *ydaS* is comparable to that of *bglG*, a well-characterized transcriptionally silent cryptic gene [^19^]. *racR* was among the most highly expressed genes in the *rac* prophage, but only to a level comparable to that of the lac repressor gene [Supplementary Figure S3].These show that YdaS is not expressed in *E.coli*, and, in light of the genetic experiments reported above, lead to the hypothesis that RacR is a repressor of this toxin.

### Binding of RacR in the IGR

RacR comprises a Helix Turn Helix (HTH) motif, and hence we investigated if it binds to DNA. The 123 bp IGR between *racR* and *ydaS* contains three slightly variant repeats of “GCCTAA” and its inverse “TTAGGC” [Figure 3B]. This is similar to the regulatory region of lambda phage, which is bound by CI and Cro, even though the exact sequences bound by the proteins are different.

To test for the binding of RacR to the IGR, we first performed a thermal shift assay with purified RacR and various nucleic acid sequences. The thermal shift assay measures the thermal denaturation temperature of a test protein. A change in this temperature in the presence of a ligand might argue in favour of an interaction between the protein and the ligand. We found that the T_D_ of RacR increased by ~5°C in the presence of *rac*R-IGR-*ydaS* or in that of a 189 bp sequence upstream of *ydaS* and including the IGR [Figure 3C]. The extended 189 bp region, including a portion of the *racR* gene, was chosen for this experiment because this included an additional half-site of the above-mentioned palindrome.

We then performed a chromatin IP of RacR::3xFLAG to test for the binding of RacR to the IGR *in vivo*. By performing qPCR against the DNA thus recovered, we found that the IGR was 2.5 fold enriched in comparison to a random region [Supplementary Figure S4-A]. Finally, we performed Electrophoretic Mobility Shift Assay (EMSA) to investigate the binding of purified RacR to the IGR. RacR formed three distinct complexes in the presence of the IGR [Figure 3D]. EMSA with a 49-bp DNA upstream of *ydaS*, containing a single copy of the repeat, also showed binding to RacR [Supplementary Figure S4-B]. Consistent with the view that the three palindromic repeats might be the sites to which RacR binds, we found only a single protein DNA complex with the 49-bp segment of the IGR. Thus, we show binding of RacR to the intergenic region between *racR* and *ydaS* both *in vitro* and *in vivo*.

### Transcriptional repression of *ydaS* is mediated by RacR binding to the IGR

Finally, to test whether the binding of RacR represses *ydaS*, we cloned the IGR upstream of *gfp-mut2* in pUA66. We monitored the promoter activity of pUA66::IGR-*gfp-mut2* in *ΔydaS-T* and in *ΔracR-ΔydaS-T* for 25 hours. We observed that the *ydaS* promoter is active only in *ΔracR-ΔydaS-T;* no fluorescence from *gfp-mut2* could be detected in *ΔydaS-T* [Figure 4A].

**Figure4:**
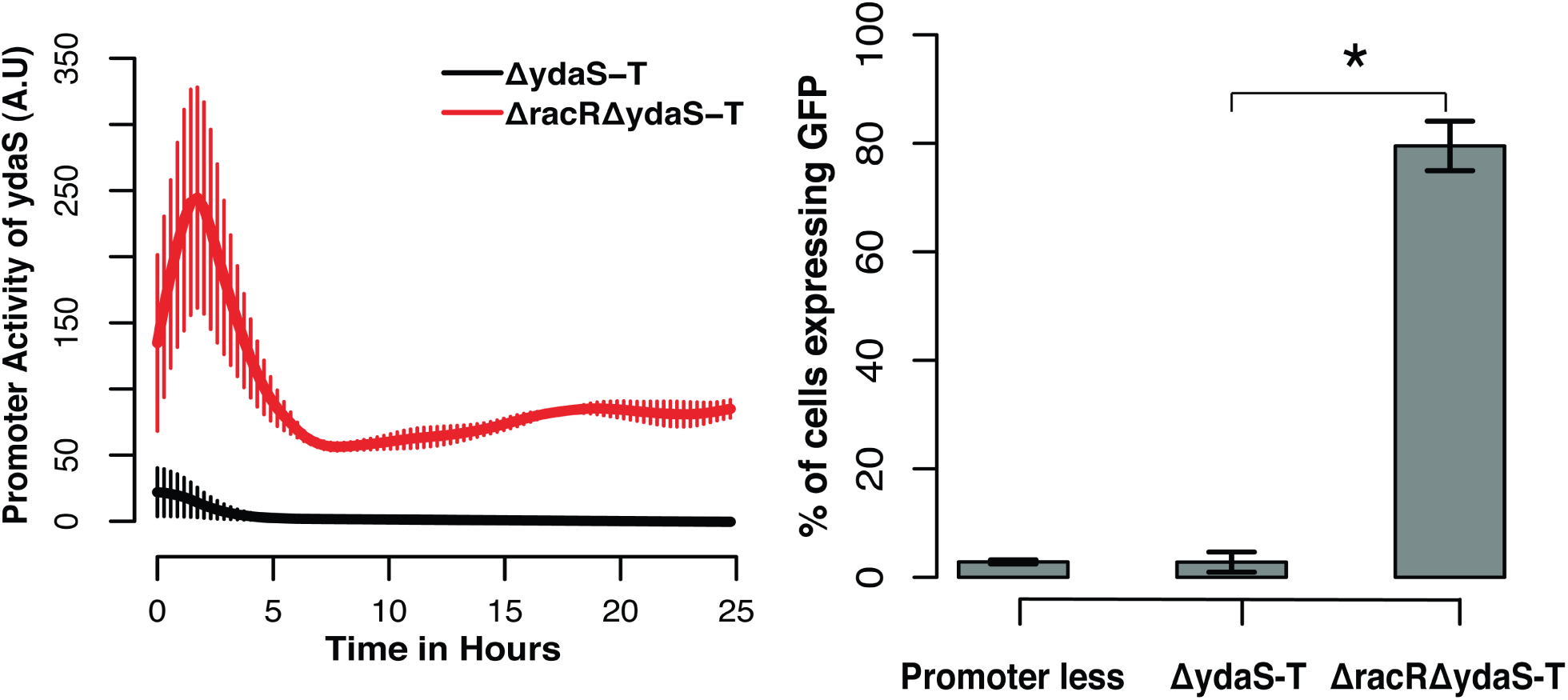
*A)IGR cloned in the low copy pUA66 plasmid, showing promoter activity of ydaS in ΔydaS-T (black line) and in ΔracRΔydaS-T (red line) strains. B) Bar graph representing the percentage of cells showing ydaS promoter Activity.* corresponds to p-value < 0.01; t-test. Error bar here represents the standard deviation computed from three independent trials.*

The maximal *ydaS* promoter activity was observed in the log phase (OD_600_ ~ 0.2-0.3). We tested the expression of *gfp-mut2* from these strains grown to mid-exponential phase using FACS. The distribution of fluorescence from pUA66::IGR-*gfp-mut2* in Δ*ydaS-T* was similar to that of the promoterless control where most cells were GFP negative. In contrast, in Δ*racR*-Δ*ydaS-T*, nearly 80% of the cells were GFP positive [Figure 4b]. Thus single copy availability of RacR from the chromosome appears to be sufficient to suppress the activity of the *ydaS* promoter from a multicopy (N = 3-4) plasmid.Thus, RacR represses transcription of *ydaS*.

## Discussion

We have shown that the expression of *ydaS* and *ydaS-T* is lethal, and we attribute the essentiality of *racR* to its role in repressing the expression of this toxin. Earlier studies have shown the presence of two toxins - KilR and RalR - in the rac prophage [^14^][^17^]. In the present work, we suggest that that YdaS-T is yet another toxin encoded by the *rac* prophage. We do not know how this toxin effects cell killing, and whether other genes in the operon to which *ydaS* and *ydaT* belong contribute to cell killing.

RacR is a repressor of *ydaS-T*, and this module is an example of a toxin-repressor system. In general, essential transcription factors are rare in *E.coli*. The essentiality of RacR is purely by virtue of its role in keeping a toxin transcriptionally silent. RacR is unlikely to have too many additional targets, because its expression level - based on RNA-seq - is very similar to that of the highly specific Lac repressor.

Among the few essential transcription factors in *E.coli* is the anti-toxin MazE. *mazE* and *mazF* are encoded on the same operon, unlike *racR-ydaS*, which make a divergent gene pair. The anti-toxin activity of MazE is primarily by protein-protein interactions with the toxin MazF. In fact, the binding of MazE to the DNA is enhanced when in complex with MazF[^20^]. Yet another essential transcription factor is the antitoxin MqsA, which again sequesters its cognate toxin MqsR. Unlike the conditional cooperativity displayed by MazE and MazF in binding to the DNA, the high stability of the MqsR-A complex makes the protein-protein interaction mutually exclusive of MqsA-DNA interactions[^21^]. In both these cases, it is apparent that the activity of the transcriptional repressor does not entirely prevent the expression of the toxin. In our case however, we could not detect the presence of YdaS protein, the expression level of the *ydaS* transcript is comparable to that of a bonafide cryptic gene across tens of RNA-seq datasets, and in the presence of RacR we cannot detect any activity from the *ydaS* promoter fused to *gfp-mut2*. The expression of YdaS-T is toxic, independent of the presence of RacR (wildtype vs. *Δrac),* which argues against the possibility of RacR interacting physically with YdaS / YdaT in suppressing its activity.

It is arguable whether RacR-YdaST can be called a toxin-antitoxin system, because the fact that the activity of the toxin is totally suppressed at the level of transcription initiation itself might render post-segregational killing downstream of the loss of the module impossible.

We propose that RacR could be functionally similar to the CI repressor of lambda prophage. The *rac* prophage has lost many of its structural genes when compared to the lambda phage [Supplementary Fig S5]. However, the organization of regulatory elements in the *rac* prophage [Figure 3B] is similar to the cl-Cro switch of lambda prophage [^22^]. There are three repeat elements in the IGR, which might be the operator of this prophage. Our observation on the formation of three distinct DNA-Protein complexes of the 123 bp IGR with increasing concentrations of RacR, suggests that the IGR might act as a complex regulatory switch that resembles the regulatory region of cI-cro of lambdoid phages [^23^].

## Materials and Methods

### Media, Strains and Plasmid Construction

*E.coli* K12 MG1655 from CGSC was used and grown at 37°C in Luria Broth (LB) or LB Agar (HiMedia). The antibiotic resistant strains were grown in antibiotics wherever required; ampicillin (100 μg/mL), kanamycin (50 μg/mL) or chloramphenicol (30 μg/mL) were used. All the knock out strains were constructed by using the one step inactivation method as described by Datsenko *et al.* using pKD13 as the template plasmid for the kanamycin resistance cassette amplification [^24^]. Tagging of *racR* with 3xFLAG at the C terminal end was done using the pSUB11 plasmid [^25^]. Ectopic expression of *racR, ydaS, ydaT* and *ydaST* were achieved by cloning them between EcoRI and SalI site of pBAD18; this brings the genes under the arabinose inducible *araBAD* promoter. The plasmid for the promoter activity was constructed by cloning the IGR in the low copy vector pUA66 between XhoI and BamHI sites. The list of strains and plasmids used in the current study is given in Supplementary Table1 and the primers used for gene deletion, validation and cloning are listed in Supplementary Table 2.

### Growth Curve and Spotting Assay

Growth curve was monitored in a 96 well plate with the final volume of 200μl using Tecan F200 reader. Overnight culture was inoculated in the ratio of 1:100 and allowed to grow till 0.4 OD. This was further diluted in fresh medium to 0.01 OD with or without 0.1% L-arabinose and A_600_ was recorded for 14 hours. For the spotting assay, appropriate overnight cultures were inoculated in LB broth containing 100 μg/mL ampicillin at 1:100 dilution, with or without 0.2% L-arabinose. The cells were collected after 5 hours and 14 hours of inoculation, serially diluted and spotted on LB agar plates containing ampicillin without arabinose.

### FACS

Overnight culture of the respective strains were inoculated in LB broth at 1:100 dilution with or without 0.2% L-arabinose. Samples were collected after 5 hours of induction, pelleted down, washed and resuspended in 500μl of saline (0.9% sodium chloride w/v). Exponentially growing cells were used as live-cell control and cells subjected to 80°C for 10 minutes were used as dead-cell control. Propidium Iodide (PI) solution (5μl of 1 mg/ml) was added to all the vials 10 minutes before acquisition of data in BD FACS Calibur. Around 20,000 cells were acquired for each sample using 488 nm excitation laser and the emission was recorded from FL2 channel that uses 585/42 BP filter, to collect the PI intensity. Intermediate population in this study is described as cells that fall between the region of live unstained control and dead control.

Exponential culture of Δ*ydaS-T* and Δ*racR*-Δ*ydaS-T* containing pUA66::IGR-*gfp*-*mut2* were pelleted, washed and resuspended in saline. GFP intensity was monitored using FL1 channel that uses 530/30 BP filter. Strain containing empty pUA66::*gfp-mut2* was used to set the background fluorescence and GFP intensity above this background was marked as positive. Data was analyzed using Flowing software (www.flowingsoftware.com/).

### Bi-Directional Search for orthologous genes

Genomes of 154 completely sequenced *E.coli* strains were downloaded from NCBI refseq ftp site. A bi-directional search for orthologous genes of the *rac* prophage, excluding pseudogenes, was performed using phmmer (Version 3.1). The E-value threshold used was 10^−20^. An ortholog presence-absence matrix was hierarchically clustered based on Euclidean distance with centroid linkage. Clustering was done using Cluster3 (bonsai.hgc.jp/~mdehoon/software/cluster/) and the heat map was generated using matrix2png (http://www.chibi.ubc.ca/matrix2png/).

### RNA - Seq Data Analysis

Raw reads from 15 different RNA-seq studies (with total of 61 fastq files) were obtained either in-house or from the NCBI GEO, or the EBI Array Express databases [Supplementary Table 3]. The SRA files from GEO were converted to fastq using fastq dump. Reads from the fastq file were aligned to NC_000913.3 genome using bwa. The aligned files were sorted using sam tools. Further, these sam files were used to get read counts per nucleotide, from which read counts per gene was generated. RPKM (Reads per Kilobase of transcript per Million mapped reads) was calculated by normalizing the raw read counts to the length of the gene and further by the total number of mapped reads for each fastq file. The distribution of RPKM values of the *rac* prophage genes were plotted as a boxplot, along with those of the *bgl* operon genes and *lacI* as reference. Because differential expression was not a goal of this study, more state-of-the-art normalization methods such as those used by EdgeR or DEseq were not required.

### Western Blotting

Total protein of *E.coli*-K12 cells was prepared and quantified using BCA Assay and 20μg of total protein was loaded in 15% SDS polyacrylamide gel. The gel was subjected to electrophoresis at 120V for 1 hour and proteins were transferred to a nitrocellulose membrane. Monoclonal anti-FLAG antibody (Sigma) was used to bind the specific protein to which the FLAG is tagged, and the signal was detected using (HRP) Horse Radish Peroxidase conjugated anti-mouse IgG. HRP luminescence was further detected by West Dura reagent (Thermo scientific). Digital images of the blots were obtained using an LAS-3000 Fuji Imager.

### Chromatin Immuno Precipitation

Immuno precipitation was done as described by Kahramanoglou et al. [^26^] except that cell lysis and DNA shearing were coupled together using Bioruptor (Diagenode) with 35 cycles (30 seconds ON and 30 seconds OFF) at high setting. Immuno precipitated samples were quantified with specific primers for the 123 bp intergenic region (IGR) and a random primer (wza), which is not the part of the *rac* prophage, using quantitative PCR. The fold enrichment was calculated using 2^−(ΔΔCt)^ as described by Mukhopadhyay et al.[^27^].

### RacR purification

RacR was cloned between the NdeI and XhoI restriction sites in a pET28a expression vector with the C-Terminal His tag. After confirmation of its sequence and orientation, this plasmid was transformed in the expression strain C41(DE3). A single colony of the C41 strain containing the pET28a::*racR* plasmid was inoculated in 5 mL LB containing 100 μg/mL ampicillin. This overnight culture was diluted to 1:100 ratio in 10 mL of fresh LB for raising the secondary inoculum. When the secondary culture reached 0.4 OD, it was seeded in fresh 1L LB in a 3L baffled flask at 37°C. When the culture reached 0.6 OD, RacR expression was induced by adding IPTG at the final concentration of 100 μM and the flask was incubated at 25°C for 5 hours. The culture was harvested and the cells were resuspended in 100 mL of Lysis buffer (50 mM Tris-pH 8.5, 500 mM Nacl, 5% Glycerol, 1% NP-40, 1x Sigma Protease Inhibitor Cocktail). The resuspended cells were sonicated for 30 cycles (30 seconds ON and 30 seconds OFF). Further, the lysate was passed through equilibrated 1 mL pre-packed Histrap column (Invitrogen) at a flow rate of 0.5mL/minute. Then the column was washed with 50 mL of elution buffer (50 mM Tris- pH 8.5, 500 mM Nacl, 5% Glycerol) containing 10 mM imidazole, and then with 20 mL of elution buffer containing 50 mM imidazole and 100 mM imidazole respectively. Finally, RacR was eluted with 10 mL of elution buffer containing 250 mM imidazole. Purified RacR was further passed through a Superdex 200 10/300 size exclusion column, which was pre equilibrated with the same elution buffer without imidazole.

### Thermal Shift Assay

0.3μM of DNA (*ydaS* with 189 bp upstream of it including a portion of *racR*, *racR-IGR-ydaS* or random DNA) was mixed with 3μM of purified RacR in the presence of 20x Sypro^®^ Orange (Sigma Aldrich), and the final volume of the reaction was adjusted to 20 μL with RacR elution buffer. Three replicates of each sample were loaded in a 384 well plate and sealed with optical adhesive cover. The fluorescence spectrum in 635 nm - 640 nm bin was recorded using ABI Via7 PCR with the standard melt curve experiment setting in which the temperature ranged from 20°C to 95°C at the rate of 1°C per minute. Denaturation temperature (T_D_) was reported as the temperature at which the maximum dF/dT was recorded, where dF/dT is the rate of change in Sypro^®^ Orange fluorescence with respect to the temperature. The data was processed and plotted using a custom R script to calculate dF/dT.

### Electrophoretic Mobility Shift Assay

The entire 123 bp IGR was PCR amplified and gel purified. Polyacrylamide gel of 6% was prepared from 40% acrylamide:bisacrylamide (80:1) stock and allowed to polymerize for 2 hours. The gel was pre-run for 30 minutes at 70 V and the wells were washed before sample loading. 20 nM of DNA was mixed with increasing concentration of RacR in 10x binding buffer (100 mM Tris Buffer-pH 8, 10 mM EDTA,1M Nacl,1mM DTT, 50% Glycerol, 0.1 mg/mL BSA) with 20 μL final volume in 0.2mL PCR tubes. These tubes were incubated at room temperature for 1 hour. After incubation, samples were mixed with 2.2 μL of 10x loading dye (10 mM Tris-pH 8, 1 mM EDTA, 50% glycerol, 0.001% bromophenol blue, 0.001% xylene cyanol.) and run at 70 V in room temperature for 90 minutes. The gel was stained using SyBr^®^ Green (Thermo Scientific) for 15 minutes. The stained gel was washed in distilled water twice and imaged using a Lab India Geldoc system.

### Promoter Activity

Promoter activity of the *ydaS* IGR was monitored by transforming the pUA66::IGR-gfp-*mut2* construct in Δ*ydaS-T* and in Δ*racR*-Δ*ydaS-T*. M9 Media with 0.2% glucose was used to culture the strains. Overnight culture containing the plasmid in the respective background strain was inoculated in the ratio of 1:100 in a 96 well flat transparent black plate (corning) with total volume of 200μL. The optical density (OD 600 nm) and the GFP intensity (excitation at 485 nm and emission at 510 nm) were measured using the Tecan multimode reader at every 16 minutes interval with continuous shaking in between at 37°C. The background optical density is subtracted by using the optical density obtained from the blank well. The background fluorescence intensity was subtracted by using the intensity obtained from the strain that has promoterless empty vector. Promoter Activity was calculated as rate of change in the GFP intensity normalized by the average OD for the given time point. PA = (smoothed) dGFP/dt/(smoothed)(OD1+OD2/2) [^28^]. Data processing and analysis were done using custom R script.

## Acknowledgements

R.K is supported by DST INSPIRE fellowship (DST/lNSPIRE Fellowship/2010). S.G and S.R is supported by DBT grant (BT/PR5801/INF/22/156/2012). S.K is supported by CSIR fellowship (09/860(0122)/2011-EMR-I). A.S.N.S is funded by Ramanujan fellowship (DST:SR/S2/RJN-49/2010). The authors thank the PTC of CCAMP and NCBS CIFF for technical support. We thank Aalap Mogre for providing his RNA seq analysis pipeline and Parul Singh for providing transcriptome data. We thank Dr. Ramaswamy and Dr. Shivaprasad for granting accession to use their lab instruments.

## SUPPLEMENTARY INFORMATION

**Supplementary Figure S1:**
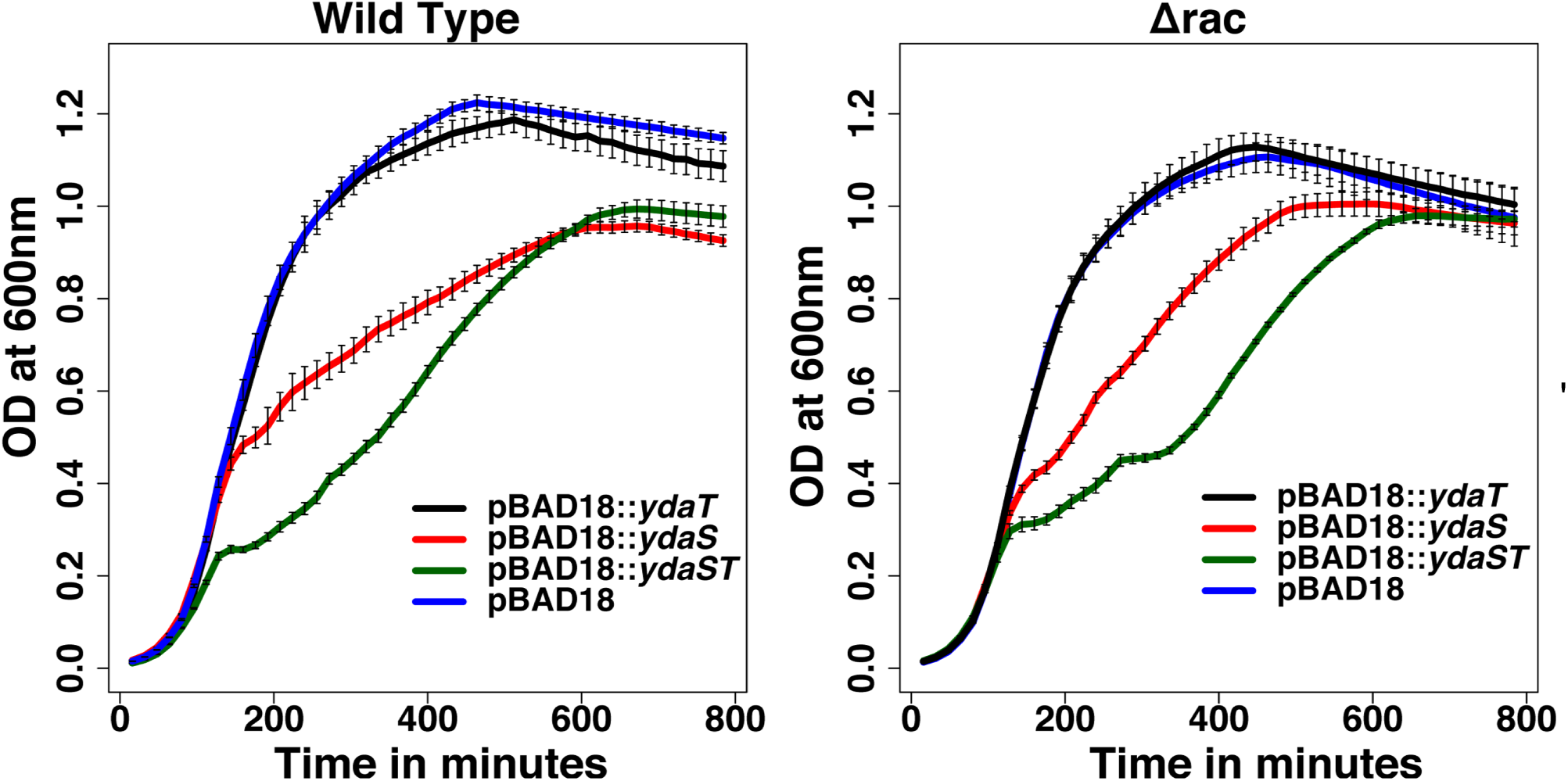
*Growth curve of the clones pBAD18-ydaS, pBAD18-ydaT, pBAD18-ydaS-T and Empty vector in wild type and in Δrac background showing that induction of ydaS and ydaS-T in tandem reduces the growth rate irrespective of the strains used*.

**Supplementary Figure S2:**
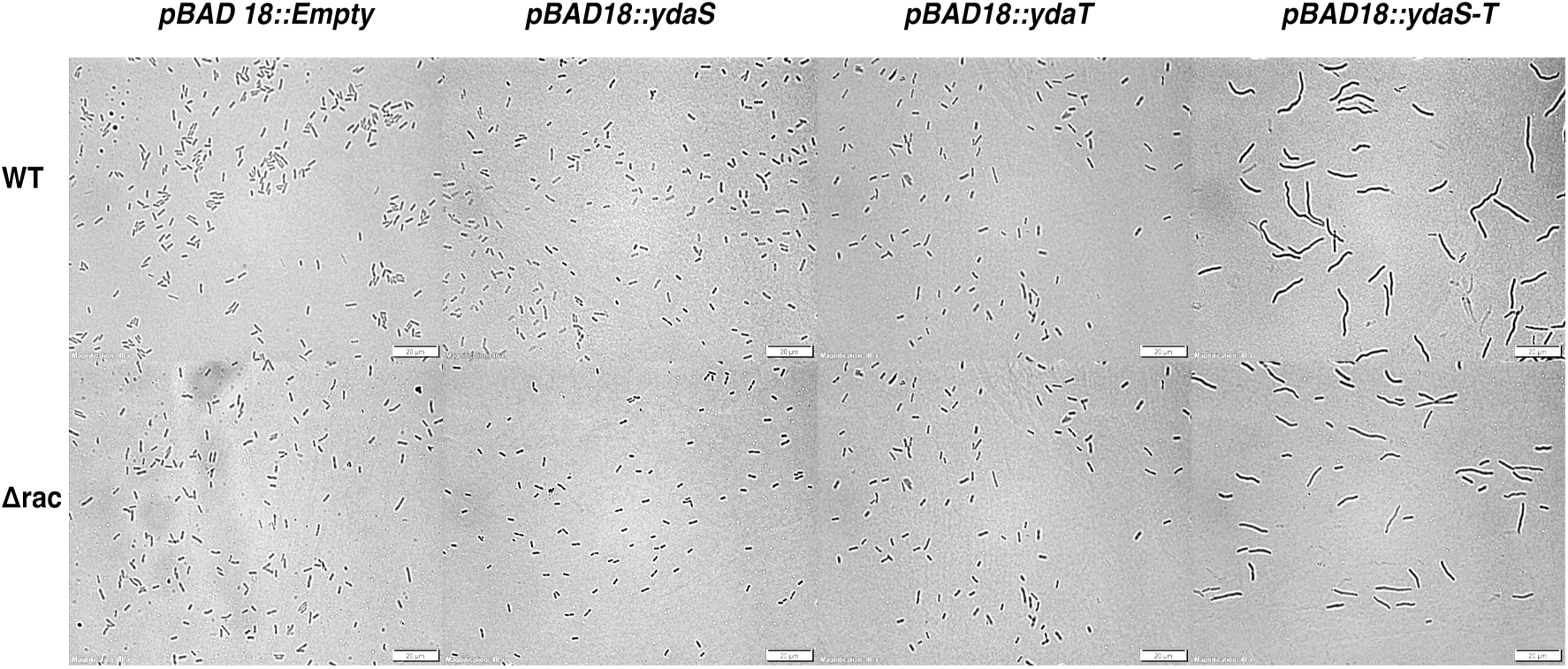
*Bright field images showing the increased cell size for the cells expressing ydaS-T when compared to the cells expressing ydaS or ydaT alone.Scale bar represents 20μm.*

**Supplementary Figure S3:**
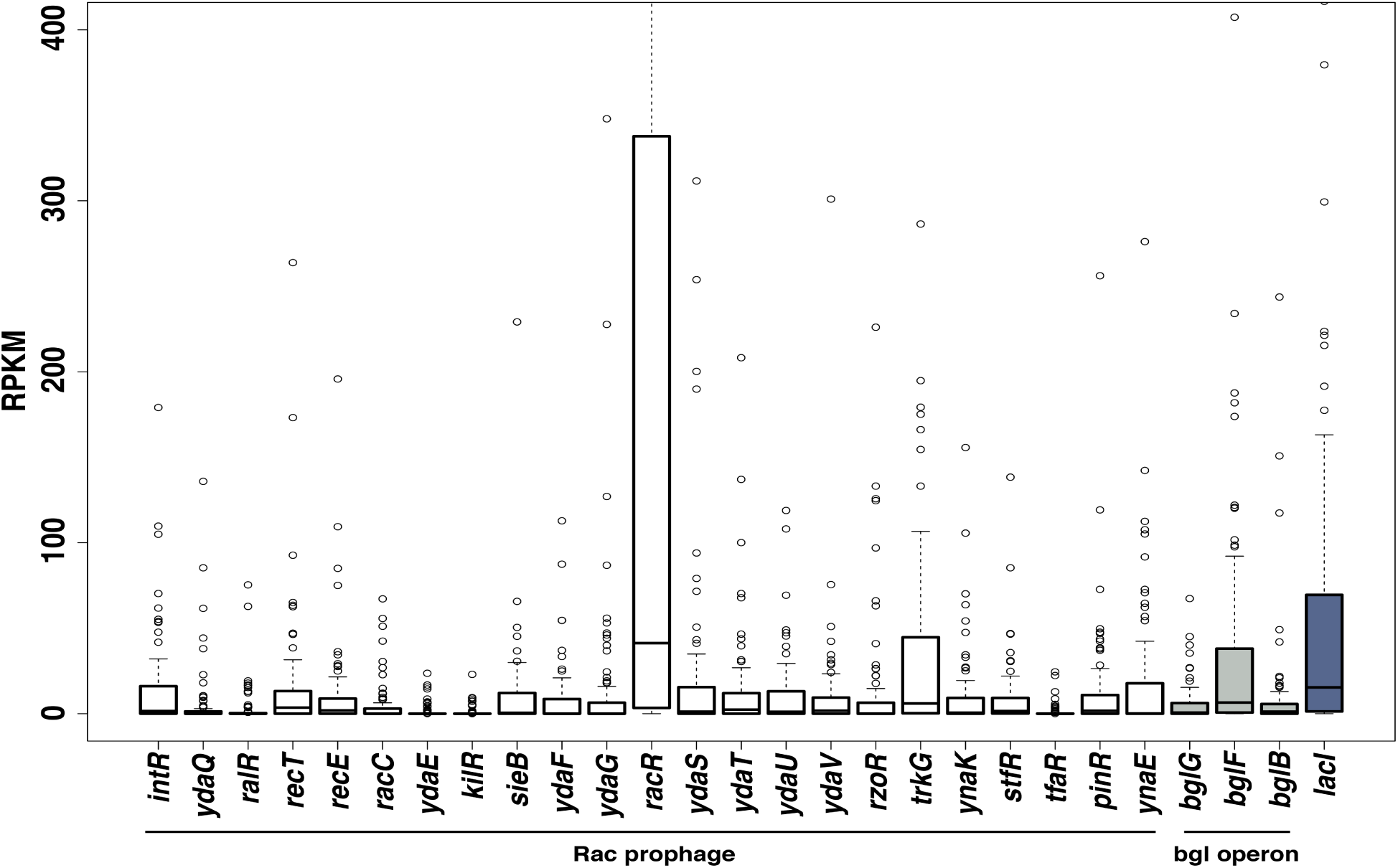
*Gene expression distribution of all the rac prophage genes is plotted as Reads Per Kilobase of transcripts per Million mapped reads (RPKM) from various RNA-Seq data. RacR is one of the few genes in the rac prophage found to be expressed across the conditions. YdaS and other toxins in the prophage is kept silent in par with the bgl operon genes.*

**Supplementary Figure S4:**
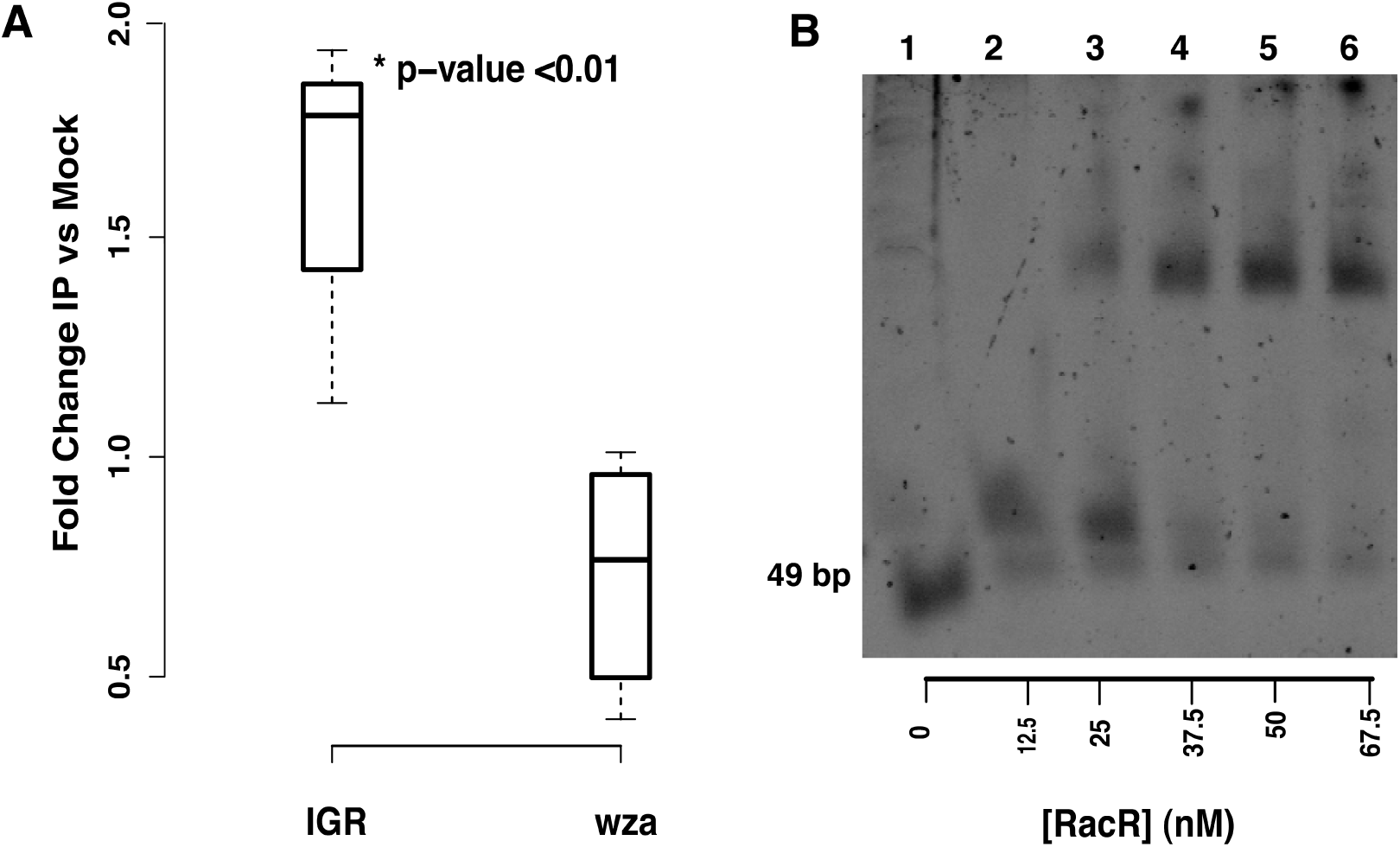
*A) Chip q-PCR showing the enrichment for IGR in the Immuno Precipitated (IP′ed) DNA. Fold change was calculated by using 2-(ΔΔCt) after normalizing the IP′ed and mock Ct to the Input Ct. Results are shown for the q- PCR done in triplicates for two biological replicate of IP′ed sample. * corresponds to p < 0.01; Wilcoxon rank sum test. B) EMSA showing the binding of RacR with 49bp region upstream ydaS. 23.5nM of 49 bp DNA was titrated against increasing concentration of RacR till 67.5 nM(Lane 2-6).Lane 1 shows 49bp without RacR.*

**Supplementary Figure S5:**
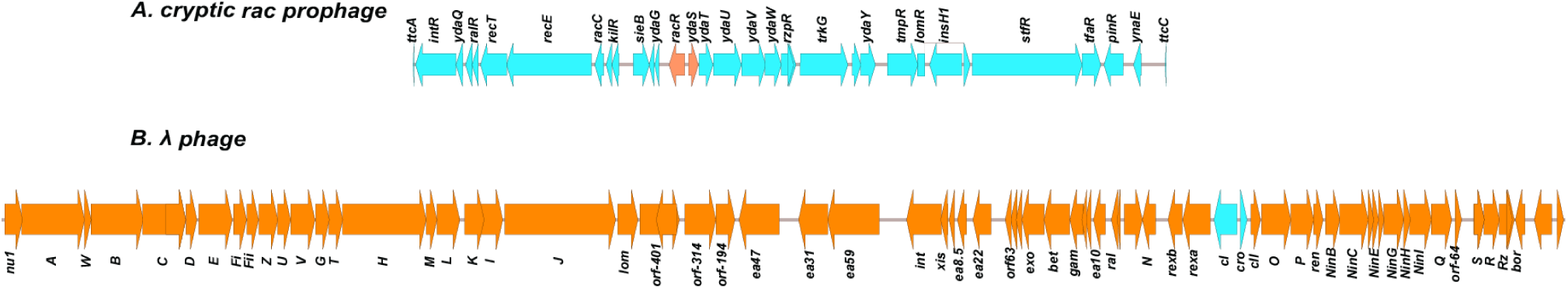
*Comparison of rac prophage with lambda prophage. Rac prophage has lost most of its structural genes when compared to the lambda prophage. The regulatory genes in both the prophages are shown in different colors. Easyfig was used to generate the map of both prophages.*

**Supplementary Table1:**
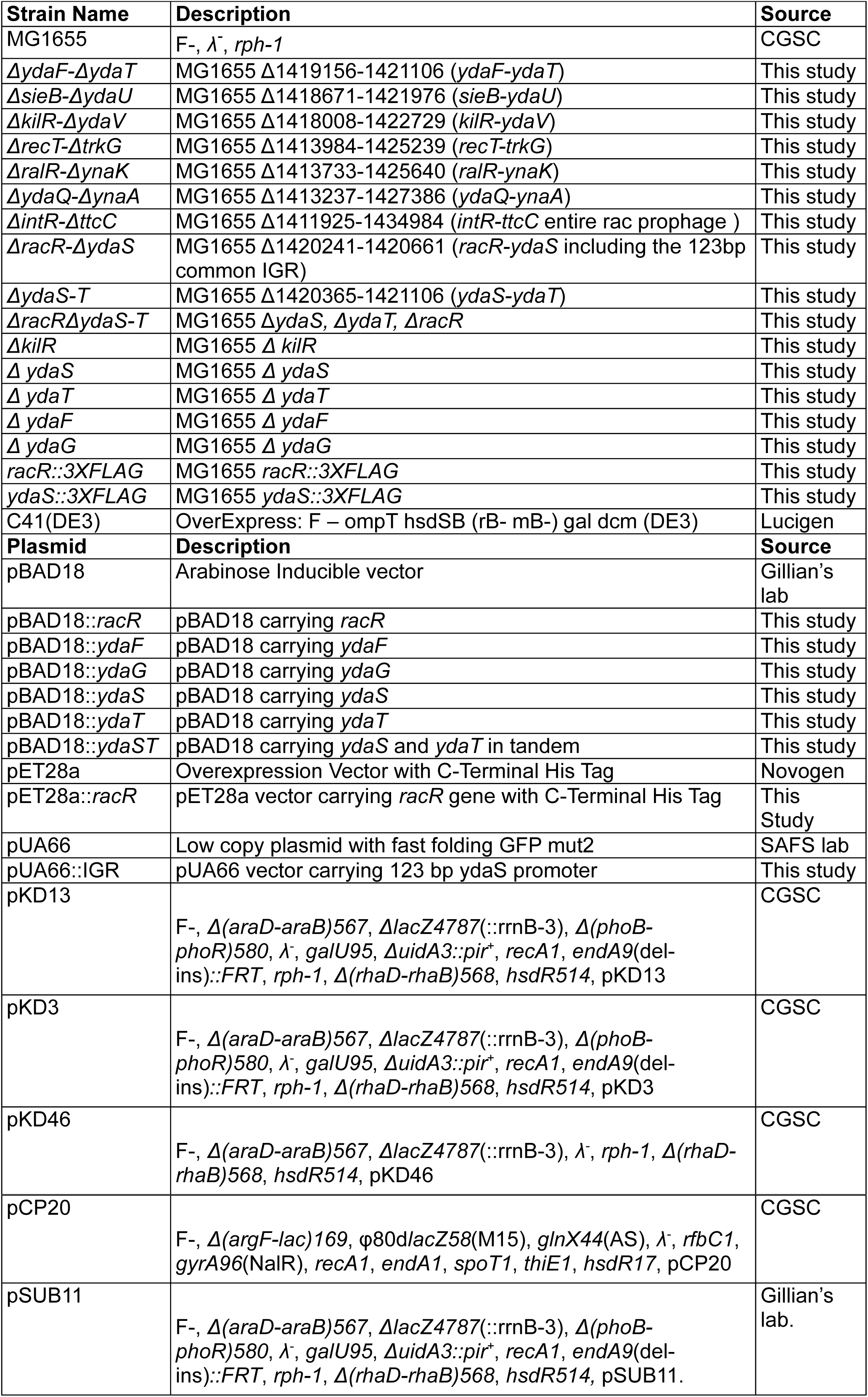
*Strains and plasmids used in this study*

**Supplementary Table2:**
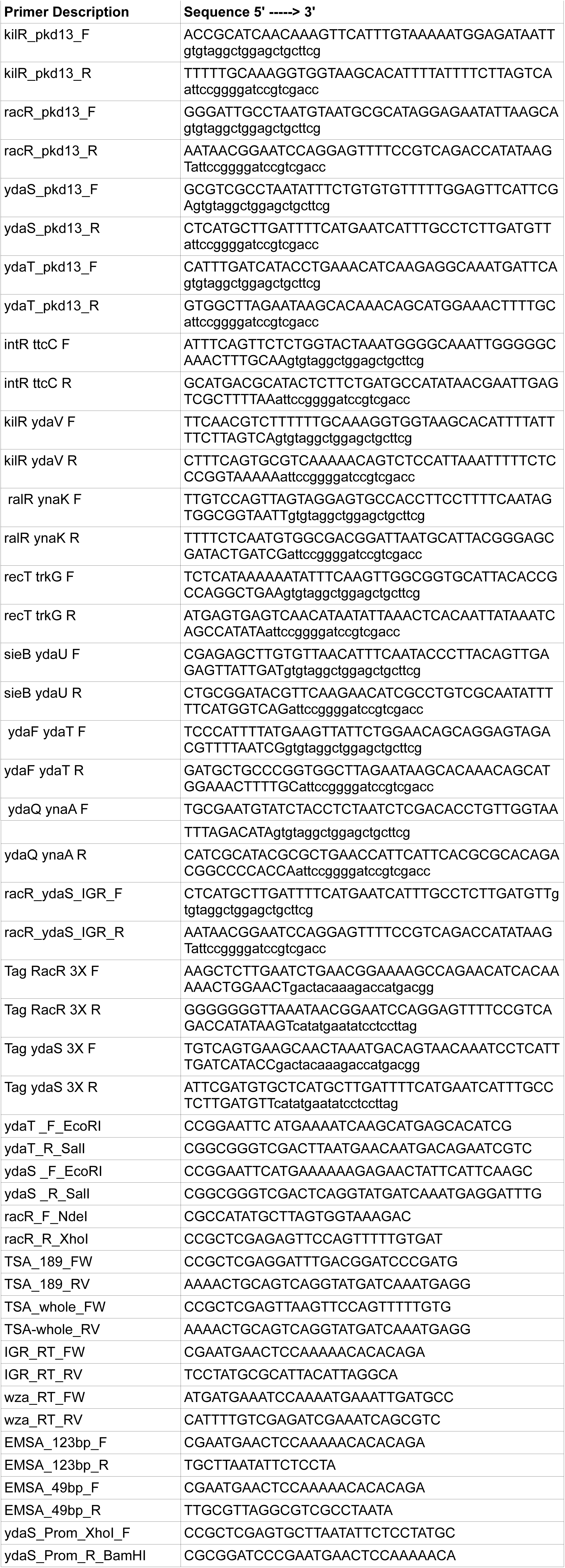
*Primers used in this study*

**Supplementary Table 3:** *List of accession numbers of RNA seq data used for calculating RPKM*

|  |  |
| --- | --- |
| GSE63858 | Ribosome profiling of *E. coli* K-12 MG1655 MOPS rich media with 0.2% glucose |
| GSE72899 | Clarifying the translational pausing landscape in bacteria by ribosome profiling [1] |
| GSE72899 | COLOMBOS v2.0: an ever expanding collection of bacterial expression compendia [2] |
| E-MTAB-2802 | Comprehensive Mapping of the *Escherichia coli* Flagellar Regulatory Network [3] |
| GSE54901 | Deciphering Fur transcriptional regulatory network highlights its complex role [4] |
| GSE66482 | Decoding genome-wide GadEWX-transcriptional regulatory networks reveals multifaceted cellular responses to acid stress in *Escherichia coli* [5] |
| GSE46740 | Genome-scale reconstruction of the sigma factor network in *Escherichia coli*: topology and functional states [6] |
| E-MTAB-2903 | Identification of bacterial sRNA regulatory targets using ribosome profiling [7] |
| GSE55199 | Global Transcriptional Start Site Mapping Using Differential RNA Sequencing Reveals Novel Antisense RNAs in *Escherichia coli* [8] |
| GSE41940 | Rho and NusG suppress pervasive antisense transcription in *Escherichia coli*[9] |
| E-MTAB-4240 | SuhB Associates with Nus Factors To Facilitate 30S Ribosome Biogenesis in *Escherichia coli*[10] |
| GSE69856 | The ribonuclease polynucleotide phosphorylase can interact with small regulatory RNAs in both protective and degradative modes [<sup>11</sup>] |
| GSE82343 | Modulation of global transcriptional regulatory networks as a strategy for increasing kanamycin resistance of EF-G mutants |
| GSE40313 | Genomic analysis reveals epistatic silencing of "expensive" genes in Escherichia coli K-12 [<sup>12</sup>] |
| In house RNA seq Data | Transcriptome data for wild type , $\Delta fis$ , $\Delta cya$ mutants in Early Exponential, Mid Exponential growth phase |

